# Transposon-insertion sequencing screens unveil requirements for EHEC growth and intestinal colonization

**DOI:** 10.1101/563007

**Authors:** Alyson R. Warr, Troy P. Hubbard, Diana Munera, Carlos J. Blondel, Pia Abel zur Wiesch, Sören Abel, Xiaoxue Wang, Brigid M Davis, Matthew K. Waldor

## Abstract

Enterohemorrhagic *Escherichia coli* O157:H7 (EHEC) is an important food-borne pathogen that colonizes the colon. Transposon-insertion sequencing (TIS) was used to identify genes required for EHEC and commensal *E. coli* K-12 growth in vitro and for EHEC growth in vivo in the infant rabbit colon. Surprisingly, many conserved loci contribute to EHEC’s but not to K-12’s growth in vitro, suggesting that gene acquisition during EHEC evolution has heightened the pathogen’s reliance on certain metabolic processes that are dispensable for K-12. There was a restrictive bottleneck for EHEC colonization of the rabbit colon, which complicated identification of EHEC genes facilitating growth in vivo. Both a refined version of an existing analytic framework as well as PCA-based analysis were used to compensate for the effects of the infection bottleneck. These analyses confirmed that the EHEC LEE-encoded type III secretion apparatus is required for growth in vivo and revealed that only a few effectors are critical for in vivo fitness. Numerous mutants not previously associated with EHEC survival/growth in vivo also appeared attenuated in vivo, and a subset of these putative in vivo fitness factors were validated. Some were found to contribute to efficient type-three secretion while others, including *tatABC, oxyR, envC, acrAB*, and *cvpA*, promote EHEC resistance to host-derived stresses encountered in vivo. *cvpA*, which is also required for intestinal growth of several other enteric pathogens, proved to be required for EHEC, *Vibrio cholerae* and *Vibrio parahaemolyticus* resistance to the bile salt deoxycholate. Collectively, our findings provide a comprehensive framework for understanding EHEC growth in the intestine.

**Author Summary:** Enterohemorrhagic *E. coli* (EHEC) are important food-borne pathogens that infect the colon. We created a highly saturated EHEC transposon library and used transposon insertion sequencing to identify the genes required for EHEC growth in vitro and in vivo in the infant rabbit colon. We found that there is a large infection bottleneck in the rabbit model of intestinal colonization, and refined two analytic approaches to facilitate rigorous identification of new EHEC genes that promote fitness in vivo. Besides the known type III secretion system, more than 200 additional genes were found to contribute to EHEC survival and/or growth within the intestine. The requirement for some of these new in vivo fitness factors was confirmed, and their contributions to infection were investigated. This set of genes should be of considerable value for future studies elucidating the processes that enable the pathogen to proliferate in vivo and for design of new therapeutics.

## Introduction

Enterohemorrhagic Escherichia coli (EHEC) is an important food-borne pathogen that causes gastrointestinal (GI) infections worldwide. EHEC is a non-invasive pathogen that colonizes the human colon and gives rise to sporadic infections as well as large outbreaks (reviewed in (1–3)). The clinical consequences of EHEC infection range from mild diarrhea to hemorrhagic colitis and include the potentially lethal hemolytic uremic syndrome (HUS) (4,5).

The paradigmatic EHEC O157:H7 strain, EDL933, caused the first recognized EHEC outbreak in 1982 (6), and it’s genome shares a common 4.1 Mb DNA backbone with the non-pathogenic laboratory strain of *E. coli* K-12, MG1655 (7–9). However, the EDL933 genome also contains 1.34Mb of chromosomal DNA that is absent from K-12, as well as a 90kb virulence plasmid pO157. EDL933-specific ‘O-islands’ encode genes recognized as the major EHEC virulence factors and are thought to have been acquired by horizontal gene transfer; many are encoded within putative prophage elements.

Although there are a variety of EHEC serotypes and the O-island complement in different EHEC isolates can differ (10), EHEC genomes all contain one or more prophages encoding Shiga toxins and the Locus of Enterocyte Effacement (LEE) pathogenicity island (11). These two horizontally acquired elements are critical EHEC virulence determinants. Shiga toxins contribute to diarrhea and the development of HUS (4,5,12). The LEE encodes a type III secretion system (T3SS) and several secreted effectors. EHEC’s T3SS mediates attachment of the pathogen to colonic enterocytes, effacement of the brush border microvilli, and the formation of actin-rich pedestal-like structures underneath attached bacteria (reviewed in (13)). Once translocated into the host cell, T3SS effectors, which are encoded both inside and outside the LEE, target diverse signaling pathways and cellular processes (14,15). A functional LEE T3SS is required for EHEC intestinal colonization in animal models as well as in humans (12,13,16–20).

In addition to the virulence factors that prompt the key symptoms of infection, EHEC also relies on bacterial factors that enable pathogen survival in and adaptation to the host environment. During colonization of the human GI tract, EHEC encounters multiple host barriers to infection, including, but not limited to stomach acid, bile, and other host- and microbiota-derived compounds with antimicrobial properties (reviewed in (21)). EHEC is known to detect intestinal cues derived from the host and the microbiota to activate expression of virulence genes and to modulate gene expression both temporally and spatially (22–25). However, comprehensive analyses of bacterial factors that contribute to EHEC survival within the host have not been reported.

The development of transposon-insertion sequencing (TIS, aka TnSeq, InSeq, TraDis, or HITS) (26–29) facilitated high-throughput and genome-scale analyses of the genetic requirements for bacterial growth in different conditions, including in animal models of infection (30–39). In this approach, the relative abundance of transposon-insertion mutants within high-density transposon-insertion libraries provides insight into loci’s contributions to bacterial fitness in different environments (40,41). Potential insertion sites for which corresponding insertion mutants are not recovered frequently correspond to regions of the genome that are required for bacterial growth (often termed “essential genes”), although the absence of a particular insertion mutant does not always reflect a critical role for the targeted locus in maintaining bacterial growth (42,43). Comparative analyses of the abundance of mutants in an initial (input) library and after growth in a selective environment (e.g., an animal host) can be used to gauge loci’s contributions to fitness in the selective condition.

Here, high-density transposon libraries were created in EHEC EDL933 and the commensal *E. coli* K-12 and used to characterize their respective in vitro growth requirements. The EHEC library was also passaged through an infant rabbit model to identify genes required for intestinal colonization. Our data indicate that during infection of the gastrointestinal tract, EHEC populations undergo a severe infection bottleneck that complicates identification of genes with true in vivo fitness defects. We used two complementary analytic approaches to circumvent the noise introduced by restrictive bottlenecks to identify genes required for colonization of the colon. More than 240 genes were found to contribute to efficient colonization of the rabbit colon. As expected, these included the LEE-encoded T3SS and *tir*, a LEE-encoded effector necessary for intestinal colonization (44). In addition, 2 non-LEE effectors and many additional new genes that encode components of the bacterium’s metabolic pathways and stress response systems were found to enable bacterial colonization of the colon. Isogenic mutants for 17 loci, including *cvpA*, a gene necessary for intestinal colonization by diverse enteric pathogens (40,45,46), were constructed, validated in the infant rabbit model and tested in vitro under stress conditions that model host-derived challenges encountered within the gastrointestinal tract. *cvpA* was found to be specifically required for resistance to the bile salt deoxycholate and therefore appears to be a previously unappreciated member of the bile-resistance repertoire of diverse enteric pathogens.

## Results and Discussion

### Identification of genes required for EHEC growth in vitro

The mariner-based Himar1 transposon, which inserts specifically at TA dinucleotides (reviewed in (47)) was used to generate a high-density transposon-insertion library in a Δ*lacI*::*aacC1* (gentamicin-resistance cassette inserted at *lacI*) derivative of EDL933. The library was characterized via high-throughput sequencing of genomic DNA flanking sites of transposon-insertion. To map the reads, we used the most recent EDL933 genome sequence (8) and annotation (NCBI, February 2017). Since this genome, unlike the initial EHEC genome (7), has not been linked to functional information (e.g., the EHEC KEGG database, KEGG reviewed in (48)), we generated a correspondence table in which the new genome annotations (RS locus tags) are linked to the original “Z numbers” (Table S1). This correspondence table enabled us to utilize historically valuable resources as well as the updated genomic sequence and should also benefit the EHEC research community. 137,805 distinct insertion mutants were identified, which corresponds to 52.5% of potential insertion mutants with an average of ∼21 reads per genotype (Fig S1A). Sensitivity analysis revealed that nearly all mutants were represented within randomly selected read pools containing ∼2 million reads. Increasing sequencing depth to ∼3 million reads had a negligible effect on library complexity, suggesting that a sequencing depth of ∼3 million reads is sufficient to identify virtually all constituent genotypes within this EHEC library (Fig S1A).

EHEC’s 6032 annotated genes were binned according to the percentage of disrupted TA sites within each gene, and the number of genes corresponding to each bin was plotted (Fig 1A). As expected for a high-density transposon-insertion library (41), this distribution was bimodal, with a minor peak comprised of genes disrupted in few potential insertion sites (Fig 1A, left), and a major peak comprised of genes that are disrupted in most or all potential insertion sites (Fig 1A, right). Based on the center of the major right-side peak, we estimate that ∼70% of non-essential insertion sites have been disrupted in this EHEC library, a degree of complexity that enabled high-resolution analysis of transposon-insertion frequency.

**Figure 1:**
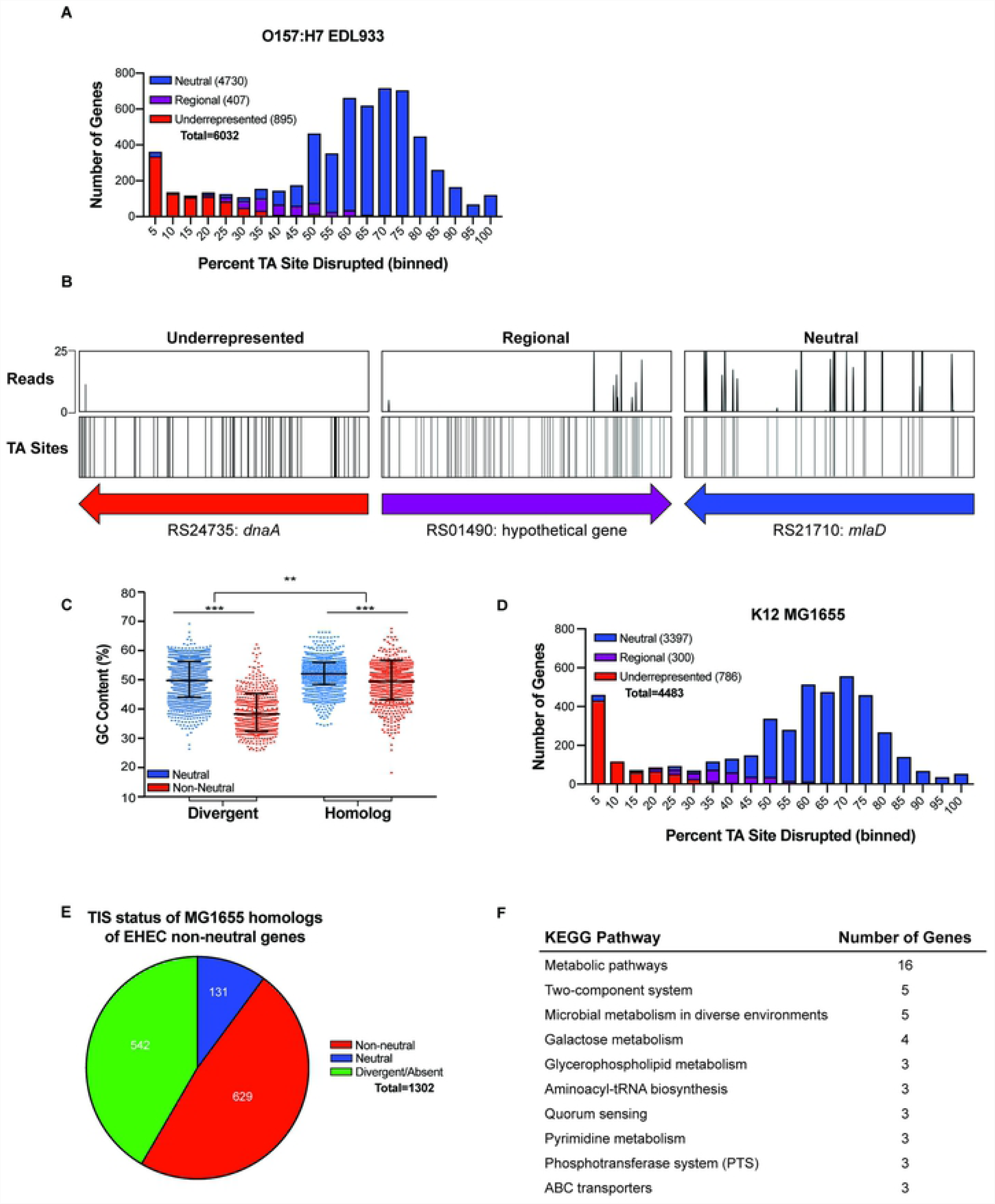
Analysis of essential genes in EHEC EDL933 and comparison to K-12 MG1655. A) Distribution of percentage TA site disruption for all genes in EHEC EDL933. Genes are classified by EL-ARTIST as either underrepresented (red), regional (purple), or neutral (blue). B) Transposon-insertion profiles of representative underrepresented, regional, and neutral genes. C) GC content (%) of EDL933 genes with and without homologs in K-12 MG1655, classified by TIS classification (neutral or non-neutral). Neutral and non-neutral genes within each gene type (divergent or homolog) are compared using a Mann-Whitney U test with a Bonferroni correction. Ratios of neutral to non-neutral genes for each gene type are compared using a Fisher’s Exact Test; (**) indicates a p-value of <0.01 and (***) indicate a p-value of <0.001. D) Distribution of percentage TA site disruption for all genes in K-12 MG1655. Genes are classified by EL-ARTIST as underrepresented (red), regional (purple), or neutral (blue) using the same parameters as for the EDL933 library. E) Genes classified as non-neutral in the ELD933 TIS library were compared to the MG1655 TIS library and categorized as either lacking a homolog (green), having the same classification in both libraries (red), or being non-neutral in EDL933 and neutral in MG1655 (blue). F) KEGG pathway information for genes that are non-neutral in EDL933 and neutral in MG1655.

Further analysis of insertion site distribution was performed using a hidden Markov model-based analysis pipeline (EL-ARTIST, see methods and (40) for detail), that classifies loci with a low frequency of transposon-insertion across the entire coding sequence as ‘underrepresented’ (often referred to as ‘essential’ genes) or across a portion of the coding sequence as ‘regional’ (Fig 1B). All other loci are classified as ‘neutral’. Of EHEC’s 6032 genes, 895 genes were classified as underrepresented, 407 as regional, and 4,730 as neutral (Fig 1A, Table S2). In general, neutral genes (blue) were disrupted in a higher percentage of TA sites than underrepresented genes (red) or regional genes (purple), which displayed low and intermediate percentage of TA site disruption, respectively (Fig 1B). Neutral genes are considered to be dispensable for growth in LB, whereas non-neutral genes (regional and underrepresented genes combined) likely have important functions for growth in this media or are otherwise refractory to transposon insertion (42,43).

We identified Z Numbers (Table S1) and the linked Clusters of Orthologous Groups (COG) (49,50) and KEGG pathways associated with the 1302 genes classified as non-neutral (underrepresented and regional). Although some loci have no COG assigned, 714 genes were assigned a COG functional category (Table S2). Each COG category was plotted against its “COG Enrichment Index”, which is calculated as the percentage of non-neutral genes in each COG category divided by the percent of the whole genome with that COG (51). A subset of COGs, particularly cell cycle control, translation, lipid and coenzyme metabolism, and cell wall biogenesis were associated with non-neutral genes at a frequency significantly higher than expected based on their genomic representation (Fig S2A). Collectively, the COG and a similar KEGG analysis (Table S3) revealed that EHEC genes with non-neutral transposon-insertion profiles are associated with pathways and processes often linked to essential genes in other organisms (52).

Non-neutral genes comprise ∼22% of EHEC’s annotated genes, a proportion of the genome that is substantially larger than the 8% and 9% observed in analogous TIS-based characterizations of *Vibrio cholerae* and *Vibrio parahaemolyticus* (40,45). Of the 1,302 non-neutral EHEC genes, only 760 are homologous to a gene in *E. coli* K-12 MG1655 (>90% nucleotide identity or >90% amino acid identity across 90+% of gene length) (Table S2, column M); thus, EDL933-specific loci comprise a high proportion (∼42%) of EHEC’s underrepresented loci. The enrichment of underrepresented loci among EHEC-specific genes, many of which were acquired by horizontal gene transfer, may reflect factors that can limit transposon-insertion other than fitness costs.

Previous analyses revealed that nucleoid binding proteins such as HNS, which binds to DNA with low GC content, can hinder Himar1 insertion (42). Consistent with this observation, genes classified as non-neutral have a lower average GC content than genes classified as neutral (Fig S2B; blue vs red distributions). The disparity in GC content between neutral and non-neutral loci is particularly marked for EHEC genes that do not have a homolog in K-12 (divergent; Fig 1C), although there is also a significant difference between the GC content of neutral and non-neutral loci with a K-12 homolog (homolog; Fig 1C). These analyses suggest that there is an association between GC content and transposon-insertion frequency in EHEC, as in other organisms, and that the prevalence of underrepresented loci among divergent loci may in part stem from the lower average GC content of these loci (Fig S2C). Additional studies are necessary to determine if the association between low GC content and reduced transposon-insertion is due to HNS-binding, other nucleoid-associated proteins, or as yet unidentified fitness-independent transposon insertion biases.

### TIS-based comparison of EHEC and *E. coli* in vitro growth requirements

To evaluate whether the abundance of non-neutral loci was specific to EHEC or was characteristic of additional *E. coli* strains, a high-density transposon-insertion library was constructed in a Δ*lacI*::*cat* (chloramphenicol-resistance cassette inserted at *lacI*) derivative of *E. coli* K-12 MG1655. EL-ARTIST analysis of the high-density K-12 library (Fig S1B) was implemented with the same parameters as those for the EHEC library analysis and classified 24% of genes as underrepresented (786 underrepresented, 300 regional and 3397 neutral; Fig 1D, Table S4). Comparison of the gene classification of homologous loci (Table S2 vs Table S4) revealed substantial concordance between the sets of genes with non-neutral insertion profiles in EHEC and K-12: 83% (629/760) of the non-neutral EHEC genes with homologs in K-12 were likewise classified as non-neutral in the *E. coli* K-12 strain (Fig 1E). Thus, analyses of non-neutral loci suggest either that the majority of ancestral loci make similar contributions to the survival and/or proliferation of EHEC and K-12 strains in LB or that they are similarly resistant to transposon-insertion.

We further explored the 131 underrepresented EHEC loci (Table S5) that were classified as neutral (able to sustain insertions) in *E. coli* K-12. Most of these genes have are linked to KEGG pathways for metabolism, particularly metabolism of galactose, glycerophospholipid, and biosynthesis of secondary metabolites (Table S5, Fig 1F). While this divergence could reflect the laboratory adaptation of the K-12 isolate, gene acquisition during EHEC evolution may have heightened the pathogen’s reliance on metabolic processes that are not critical for growth of K-12. Such ancestral genes may be useful targets for antimicrobial agents, as they might antagonize EHEC growth without disruption of closely related commensal Enterobacteriaceae populations.

### Comparison of TIS and deletion-based gene classification

The sets of genes classified as underrepresented or regional in EHEC and K-12 transposon libraries were compared to the 300 genes classified as essential in the K-12 strain BW25113 based on their absence from a comprehensive library of single gene knockouts (53–55). 98% of these genes (294/300) were also classified as underrepresented or regional in EDL933 and MG1655 (Table S2 and S4). The few loci previously classified as essential but not found to be underrepresented or regional in our analysis include several small genes, whose low number of TA sites hampers confident classification. One gene in this list, *kdsC*, was found to have insertions across the gene in both EDL933 and MG1655 (Fig S2D). *kdsC* knockouts have also been reported previously (56), confirming that this locus is not required for K-12 growth despite the absence of an associated mutant within the Keio collection. Thus, underrepresented and regional loci encompass, but are not limited to, loci previously classified as essential.

Several factors likely account for over-estimation of loci as underrepresented or regional. First, loci can be classified as underrepresented even when viable mutants are clearly present within the insertion library (Fig 1A); insertions simply need to be consistently less abundant across a segment of the gene than insertions at other (neutral) sites. Loci may also be classified as underrepresented due to fitness-independent insertion biases, as discussed above (42,43). Additional evidence that loci categorized as non-neutral by transposon-insertion studies are not necessarily essential for growth was provided by a recent study of essential genes in E. coli K-12 (57). However, the more expansive non-neutral classification can provide insight into loci that enable optimal growth, in addition to those that are required.

### Identification of EHEC genes required for growth in vivo

To identify mutants deficient in their capacity to colonize the mammalian intestine, the EHEC transposon library was orogastrically inoculated into infant rabbits, an established model host for infection studies (12,44,58,59). Transposon-insertion mutants were recovered from the colon at 2 days post-infection, and the sites and abundance of transposon-insertion mutations were determined via sequencing, as described above. The relative abundance of individual transposon-insertion mutants in the library inoculum was compared to samples independently recovered from the colons of 7 animals to identify insertion mutants that were consistently less abundant in libraries recovered from the colon. Under ideal conditions, this signature is indicative of negative selection of the mutant during infection, reflecting that the disrupted locus is necessary for optimal growth within the intestine.

Sequencing and sensitivity analyses of the 7 passaged libraries revealed that they contained substantially fewer unique insertion mutants than the library inoculum (23-38% total mutants recovered, ∼30,000 of 120,000) (Fig S1C-J). These data are suggestive of population constrictions that could have arisen from 2 distinct but not mutually exclusive causes: 1) negative selection, leading to depletion of mutants deficient at in vivo survival or intestinal colonization; and/or 2) infection bottlenecks, population constrictions that lead to stochastic reductions in the average number of insertions per gene, independent of genotype or selective pressures. We binned genes according to the percentage of TA sites disrupted within their gene sequences and plotted the number of genes corresponding to each bin for both the inoculum (Fig 2A-top) and a representative rabbit-passaged sample (Fig 2A-bottom). The passaged sample exhibited a marked leftward shift relative to the inoculum, a signature indicative of population constriction due to an infection bottleneck (41,60).

**Figure 2:**
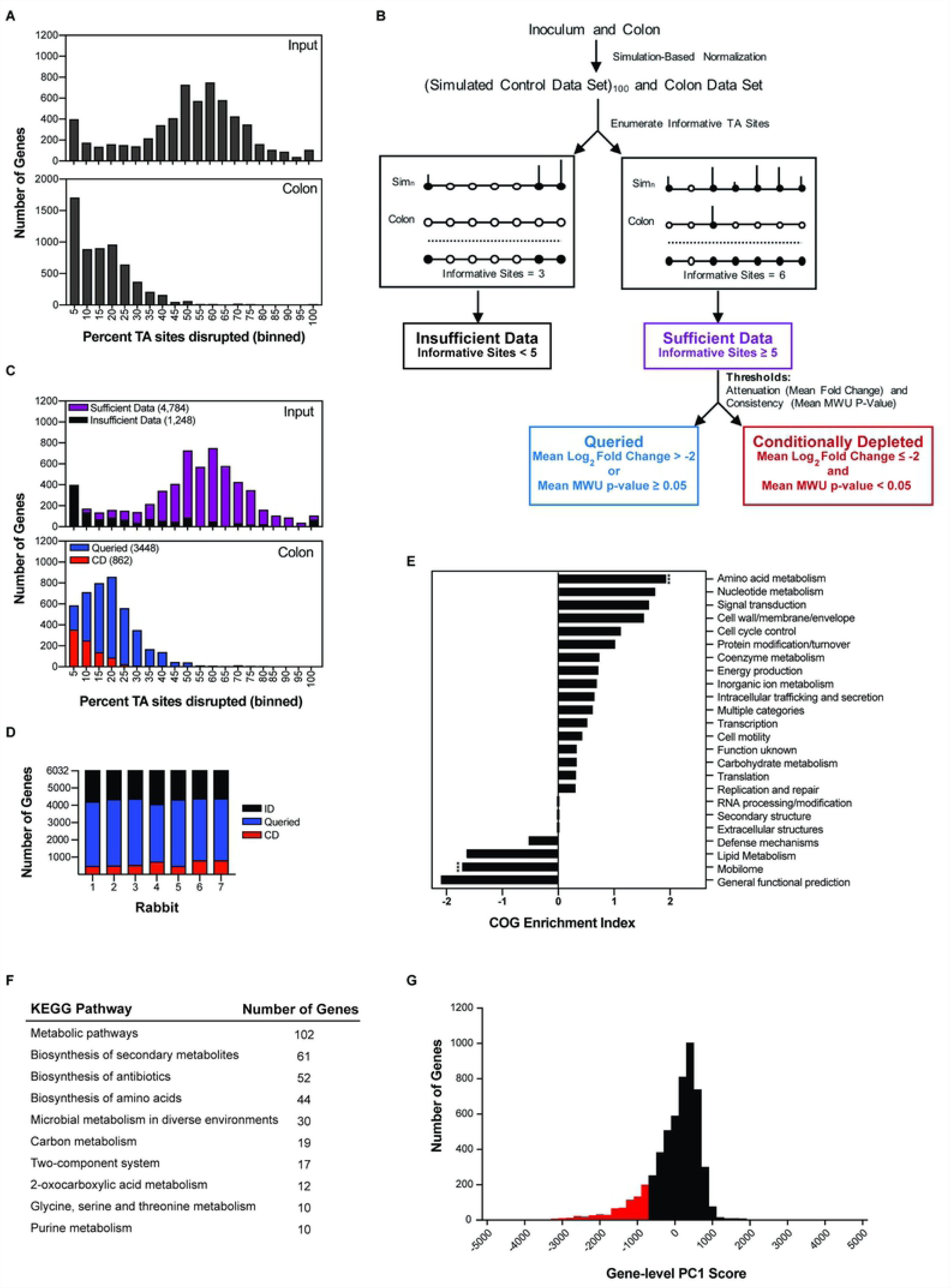
Identification of EHEC genes required for intestinal colonization. A) Distribution of percentage TA site disruption for all genes in EDL933 in the library used to inoculate infant rabbits (top) and in a representative library recovered from a rabbit colon two days after infection (bottom). B) Schematic of Con-ARTIST classification scheme. Con-ARTIST utilizes iterative resampling of the inoculum data set to generate 100 simulated control data sets and compares relative abundance of mutants in these simulated control data sets relative to the passaged library. Genes with sufficient data (≥ 5 TA sites disrupted) are then classified based on a dual standard of attenuation (mean log_2_ fold change ≤ −2) and consistency (Mann Whitney U p-value <0.05) as either queried (blue) or conditionally depleted (red). C) Distribution of percentage TA site disruption for all genes in the inoculum library (top) and a representative library recovered from the rabbit colon (bottom) overlaid with the classifications described in panel B. Genes with insufficient data are removed from the bottom panel. D) Distribution of Con-ARTIST gene classifications (insufficient data (ID, black), queried (Q, blue), conditionally depleted (CD, red)) in each library recovered from seven infant rabbit colons two days post infection as compared to the inoculum. E) Conditionally depleted genes (defined by Con-ARTIST consensus approach) by Clusters of Orthologous Groups (COG) classification. COG enrichment index (displayed as log2 enrichment) is calculated as the percentage of the CD genes assigned to a specific COG divided by the percentage of genes in that COG in the entire genome. A two-tailed Fisher’s exact test with a Bonferroni correction was used to test the null hypothesis that enrichment is independent of TIS classification. (***), p-value <0.0001. F) KEGG pathways of EHEC genes classified as conditionally depleted by Con-ARTIST consensus approach. G) Distribution of PC1 scores across all EHEC genes. Red bins fall within the lowest 10% of PC1 scores.

The TIS data was further analyzed using the Con-ARTIST pipeline. Con-ARTIST uses iterative simulation-based normalization to compensate for experimental bottlenecks to facilitate discrimination between stochastic reductions in genotype abundance and reductions attributable to bona fide negative selection (mutants for which there was a fitness cost in the host environment) (40). The Con-ARTIST analysis protocol and subsequent gene classification is schematized in Figure 2B. For libraries recovered from each rabbit, we used this workflow to classify genes as ‘conditionally depleted’ (red, CD), ‘queried’(blue) or ‘insufficient data’ (black) (Fig 2B) compared to the inoculum library. CD genes contain sufficient insertions for analysis (see methods) and meet a standard of a 4-fold reduction in read abundance that is consistent across TA sites in a gene (Fig 2B). Queried genes contained sufficient insertions for analysis but failed to meet this criterion. Genes classified as insufficient do not contain sufficient insertions for analysis. The output of gene categorization using these thresholds is displayed for a single rabbit in Fig 2C (additional animals in Fig S3A-G) and summarized for all animals in Fig 2D.

The restrictive bottleneck and animal to animal variation led to differences in the numbers of genes classified as CD in different rabbits (Fig 2D). Due to this variability, an additional criterion that genes be classified as CD in 5 or more of the 7 animals analyzed was imposed to create a consensus cutoff. In contrast to the >2000 genes classified as conditionally depleted in one or more animals, only 243 genes were classified as conditionally depleted across 5 or more animals (Fig S3H, Table S6). These relatively stringent standards were imposed in order to identify robust candidates for genes that facilitate EHEC intestinal growth, despite the limitations of the infection bottleneck in this experimental model. Therefore, we do not conclude that genes classified as “Queried” (3860) or “Insufficient Data” (1926) are *not* attenuated relative to the wild type strain in vivo; it is likely that the list of CD loci (Table S6) is incomplete.

Using our Z correspondence table (Table S1), 89% (217/243) genes classified as conditionally depleted were assigned to a COG functional category. CD genes were frequently associated with amino acid and nucleotide metabolism, signal transduction, and cell wall/envelope biogenesis, but only amino acid metabolism reached statistical significance after correction for multiple hypothesis testing (Fig 2E). These genes are also associated with KEGG metabolic pathways (particularly amino acid metabolism), several two-component systems, including *qseC*, which has previously been implicated in EHEC virulence gene regulation, and lipopolysaccharide biosynthesis (22) (Table S7, Fig 2F). 33 of the 243 CD genes are EHEC specific, whereas the remaining 210 have homologs in K-12 (Table S6), highlighting the importance of conserved metabolic pathways in the pathogen’s capacity to successfully colonize its colonic niche. Similar metabolic pathways were also found to be important for *V. cholerae* growth in the infant rabbit small intestine (40,61), and raising the possibility of targeting metabolic pathways such as those for amino acid biosynthesis with antibiotics (62–65).

The stochastic loss of individual insertion mutants in severely bottlenecked data can hinder Con-ARTIST-based identification of CD genes. In particular, meeting the pipeline’s consistency threshold as measured by the Mann Whitney U (MWU) p-value (Fig 2B) is difficult because severe bottlenecks drastically decrease the number of individual transposon-insertion mutants per gene; therefore, the mutant replicates needed to demonstrate consistency are not present. Queried genes often did not meet the MWU p-value cut-off, even though fold change information may suggest marked attenuation. To reclaim some of the mutants that were not classifiable by Con-ARTIST, we also used Comparative TIS (CompTIS), a principal components analysis (PCA)-based framework (66), to compare the 7 libraries recovered from rabbit colons. PCA is a dimensional reduction approach used to describe the sources of variation in multivariate datasets. Recently, we found that PCA is useful for identifying genes whose inactivation leads to mutant growth phenotypes that are consistent across TIS replicates (66). Here, we applied CompTIS as an alternative approach to identify genes with phenotypic consistency (inability to colonize the rabbit colon) in all 7 rabbit replicates.

To perform CompTIS, the fold change of each gene from the seven colon libraries was subjected to gene level PCA (glPCA) (see methods and (66)), with each library recovered from a rabbit colon representing a replicate. glPC1 describes most of the variation in the animals (Fig S3I) and reports a weighted average of the fold change values for each gene across the 7 animals (Table S6). The signs and magnitudes of PC1 were all similar (Fig S3J), indicating that each rabbit contributes approximately equally to PC1, as expected for biological replicates. The distribution of glPC1 scores is continuous (Fig 2G) and describes each gene’s contribution to EHEC intestinal colonization. Most genes have a glPC1 score close to zero (average PC1=0), suggesting that they do not contribute to colonization. However, the distribution includes a left tail beginning at PC1 scores of approximately −900 that encompassed the lowest 10% of scores (Fig 2G, Fig S3K), which likely correspond to genes contributing to colonization. This list of 541 genes included nearly all (85%) of the genes classified as CD by the more conservative Con-ARTIST analysis outlined above (Fig 2B). This method allows for identification of additional candidate genes required for survival/growth in vivo. For example, the PCA approach captured genes such as *ler* (PC1 = −2290), a critical activator of the LEE T3SS (67), which was classified as queried by Con-ARTIST due to the relative paucity of unique insertion mutants.

### Analyses of the requirement for T3SS and its associated effectors in colonization

To begin to assess the accuracy of our gene classifications using the Con-ARTIST consensus approach and CompTIS, we examined classifications within the LEE pathogenicity island, which encodes the EHEC T3SS and plays a critical role in intestinal colonization (12,16–19). The LEE is comprised of 40 genes, including genes encoding the structural components of the T3SS, some of the pathogen’s effectors, their chaperones, and Intimin (encoded by *eae*), the adhesin that binds to the translocated Tir protein. In infant rabbits, previous studies using single deletion mutants revealed that *tir, eae*, and *escN*, the T3SS ATPase, were all required for colonization (12,44). We observed a marked reduction in the abundance of insertions across nearly the entire LEE in the samples from the rabbit colons relative to the simulation-normalized input reads (Fig 3A). The 3 genes previously found to be required for colonization (*tir, eae* and *escN*) were classified as CD using the Con-ARTIST consensus approach, enhancing confidence in this scheme. Furthermore, 8 additional LEE-encoded genes critical for T3SS activity, including translocon T3SS components (*espB, espD*, and *espA*) and structural components (*escD, escQ, escV, escI*, and *escC*) were also classified as CD using this scheme (Fig 3AB, Table S6) (16,68–72). Our findings provide additional strong evidence that the LEE T3SS is critical for EHEC proliferation in the intestine. However, many LEE-encoded genes had insufficient data to enable classification via the Con-ARTIST consensus approach.

**Figure 3:**
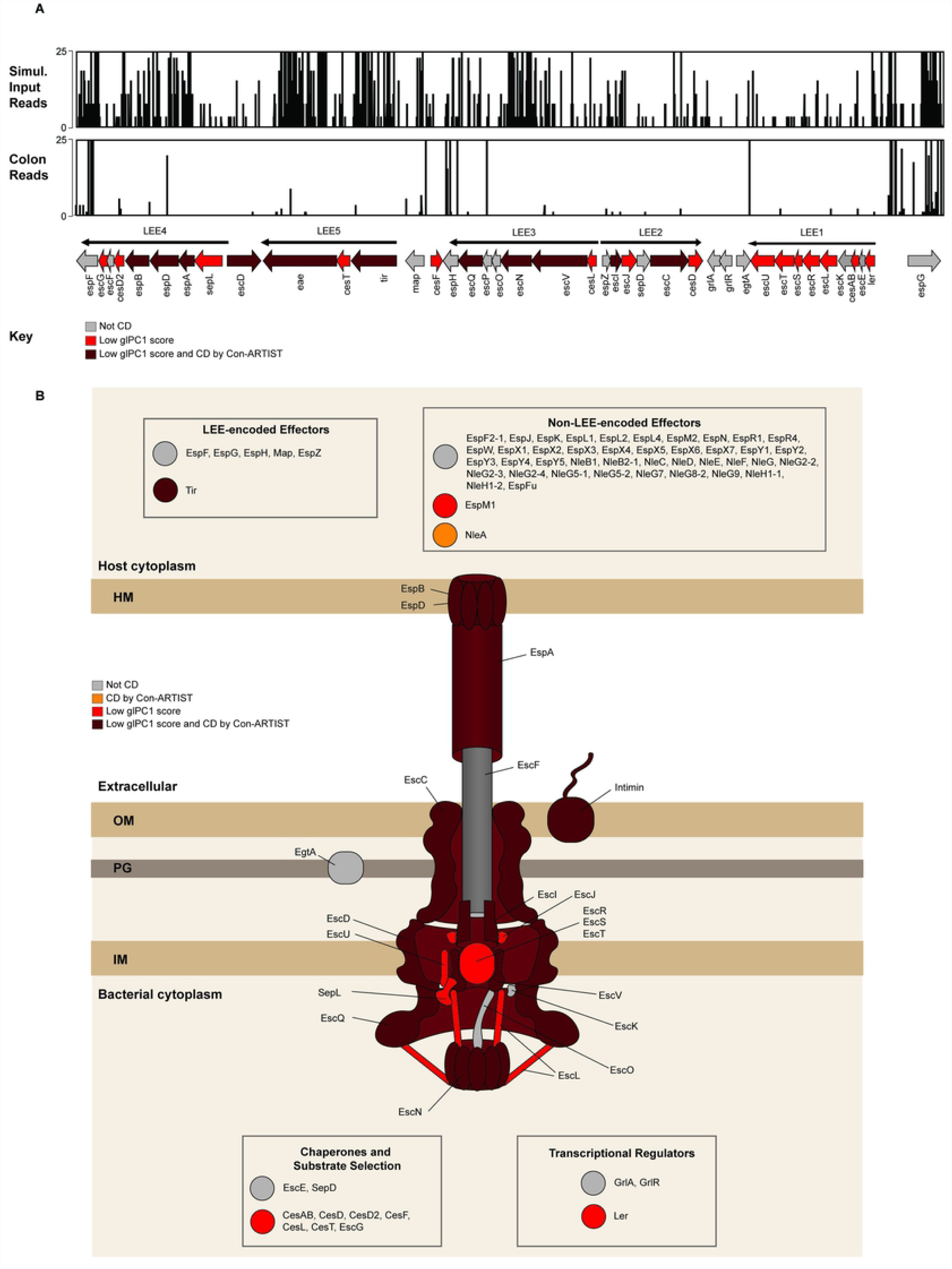
Con-ARTIST and CompTIS-based classification of LEE genes and T3SS effectors. A) Artemis plots of reads in the LEE pathogenicity island in the control-simulated inoculum library (top) and a representative library recovered from the rabbit colon (middle). The genes in the LEE are displayed at the bottom. The color of the gene corresponds to its classification. Maroon genes were categorized as conditionally depleted by Con-ARTIST consensus and fell in the bottom 10% of glPC1 scores by CompTIS; red genes had a glPC1 score in the bottom 10% of the distribution, but not classified as CD by Con-ARTIST. Gray genes did not meet the glPC1 cutoff and were not classified by Con-ARTIST. B) Schematic showing classification of the LEE genes and non-LEE-encoded effectors. Color symbols as above. Orange indicates the gene was identified as CD by Con-ARTIST and had a glPC1 score above 10%. LEE-encoded effectors, non-LEE encoded effectors, chaperones/substrate selection proteins, and regulators are indicated in boxes.

In contrast to Con-ARTIST analytic approach, with the PCA-based CompTIS analysis, we were able to assess the contribution of all of the genes in the LEE and identify several more genes likely to be important for in vivo colonization. With this approach, most LEE genes had glPC1 scores in the bottom 10% of the distribution (Table S6, Fig 3B). Notably, the genes that were not in this portion of the distribution included 4 effectors (*espF, map, espH, espG*). Previous studies in infant rabbits showed that *espG* and *map* were dispensable for colonic colonization and that *espH* and *espF* mutants only had modest colonization defects (44), lending credence to PCA-based classification.

EHEC has a large suite of non-LEE encoded effectors (Nle), many of which reside within prophage elements. Only 2 of 43 Nle genes (*nleA, espM1)* were classified as CD by Con-ARTIST or were found within the bottom 10% of glPC1 scores by CompTIS (Fig3B, Table S6), suggesting that only a small subset of EHEC effectors are critical for colonization, while other effectors likely play auxiliary roles. NleA was previously reported to be important for colonic colonization by a related enteric pathogen, *Citrobacter rodentium* (73), and is thought to suppress inflammasome activity (74), while EspM1 is thought to modulate host actin cytoskeletal dynamics (75,76). Additional studies are warranted to confirm and further explore how these 2 effectors play pivotal roles promoting intestinal colonization.

### Validation of colonization defects in non-LEE encoded genes classified as conditionally depleted

We performed further studies of 17 conditionally depleted genes/operons that had not previously been demonstrated to promote EHEC intestinal colonization. The Con-ARTIST consensus approach and CompTIS classified all of these genes as conditionally depleted except one, *hupB*, which was classified as queried by Con-ARTIST but within the bottom 10% of glPC1 scores (Fig 4A). Mutants with in-frame deletions of either single loci (*agaR, cvpA, envC, htrA, hupB, mgtA, oxyR, prc, sspA, sufI, tolC*, and RS09610, a hypothetical gene of unknown function) or operons with one or more genes classified as conditionally depleted (*acrAB, clpPX, envZompR, phoPQ, tatABC*) were generated. Then, each mutant strain was barcoded with unique sequence tags integrated into a neutral locus in order to enable multiplexed analysis. The in vitro growth of the barcoded mutants was indistinguishable from that of the WT strain (Fig S4A), suggesting that the transposon mutants’ in vivo attenuation is not explained by a generalized growth deficiency.

**Figure 4:**
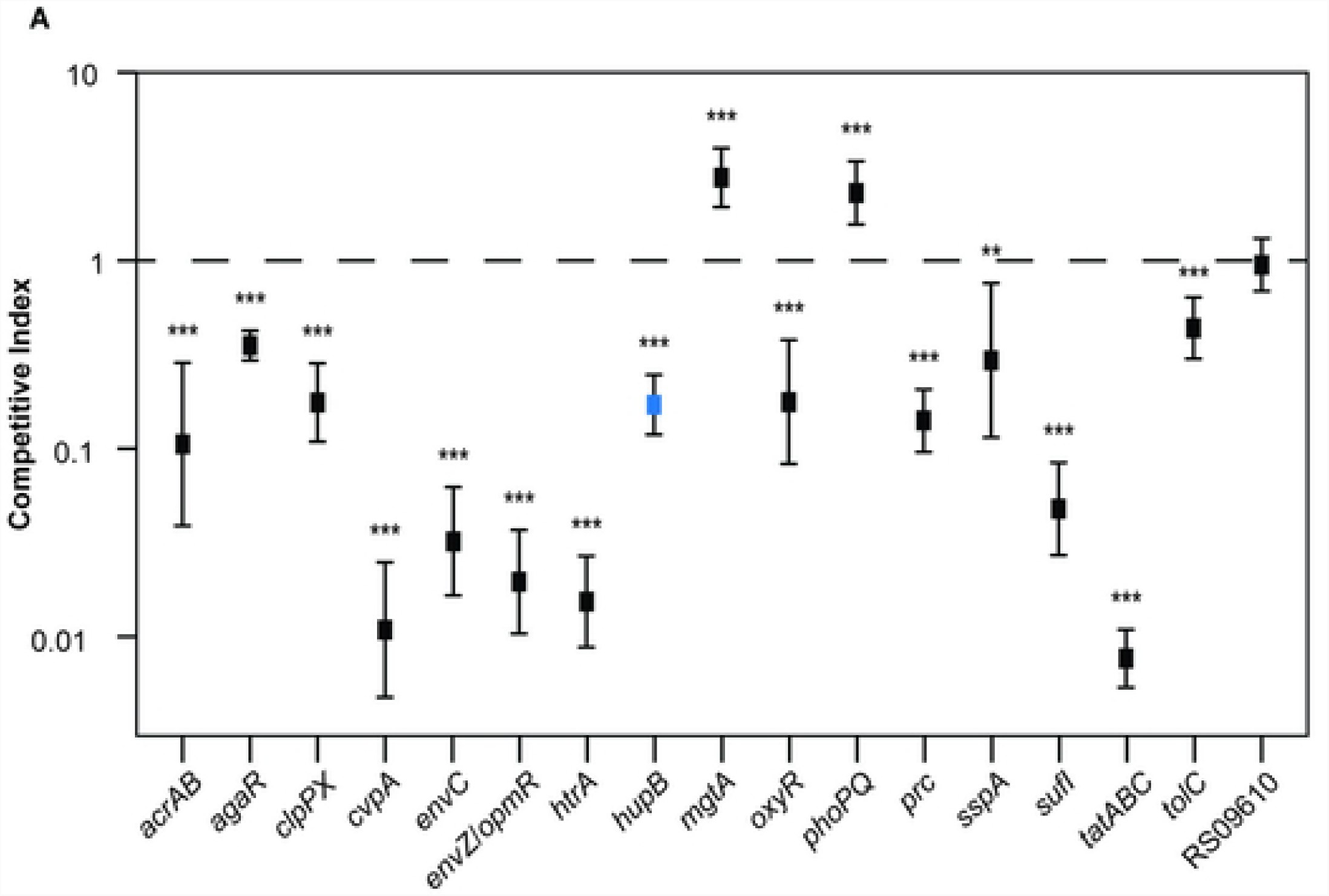
Validation of colonization defects in selected mutants. A) Competitive indices of indicated mutants vs wild type EHEC. Bar-coded mutants were co-inoculated with bar-coded wild-type EHEC into infant rabbits and recovered two days later from the colons of infected rabbits. Relative abundance of each mutant was determined by sequencing the barcodes. (**) p-values < 0.01 and (***) p-value <0.001. Δ*hupB*, which had a glPC1 score in the bottom 10%, but was not classified as CD by Con-ARTIST, is highlighted in blue.

The barcoded mutants, along with the barcoded WT EHEC, were co-inoculated into infant rabbits to compare the colonization properties of the mutants and WT. The relative frequencies of WT and mutant EHEC within CFU recovered from infected animals was enumerated by deep sequencing of barcodes, and these frequencies were used to calculate competitive indices (CI) for each mutant (i.e., relative abundance of mutant/WT tags in output normalized to input). 14 of the 17 mutants tested had CI values significantly lower than 1, validating the colonization defects inferred from the TIS data (Fig 4). In aggregate, these observations support our experimental and analytical approaches and suggest that many of the genes classified as CD by the Con-ARTIST consensus approach and/or have low PC-1 scores may also contribute to intestinal colonization.

### Many conditionally depleted loci exhibit reduced T3SS effector translocation and/or increased sensitivity to extracellular stressors

The many new genes implicated in EHEC colonization by the TIS data could contribute to the pathogen’s survival and growth in vivo by a large variety of mechanisms. Given the pivotal role of EHEC’s T3SS in intestinal colonization, as well as previous observations that factors outside the LEE can regulate T3SS gene expression and/or activity (reviewed in (67)), we assessed whether T3SS function was impaired in the 11 mutants with CIs <0.3 (Fig 4). Translocation of EspF (an effector protein) fused to a TEM-1 beta-lactamase reporter into HeLa cells was used as an indicator of T3SS functionality (77). An Δ*escN* mutant, which lacks the ATPase required for T3SS function, was used as a negative control.

Deletions in three protease-encoded genes, *clpPX, htrA*, and *prc*, were associated with reduced EspF translocation (Fig 5A). Both ClpXP and HtrA have been implicated in T3SS expression/activity in previous reports (78–81). The ClpXP protease controls LEE gene expression indirectly by degrading LEE-regulating proteins RpoS and GrlR (82). The periplasmic protease HtrA (aka DegP) has been implicated in post-translational regulation of T3SS as part of the Cpx-envelope stress response (80,81). Interestingly, *prc*, which also encodes a periplasmic protease (82), also appears required for robust EspF translocation. Prc has been implicated in the maintenance of cell envelope integrity under low and high salt conditions in *E. coli* K-12 (83). Consistent with this observation, in high osmolarity media a Δ*prc* EHEC mutant exhibited cell shape defects (Fig S4B). Deficiencies in the cell envelope associated with absence of Prc may impair T3SS assembly and/or function, perhaps also by triggering the Cpx-envelope stress response. Together, these observations suggest that in vivo these three proteases modulate T3SS expression/function, thereby promoting EHEC intestinal colonization.

**Figure 5:**
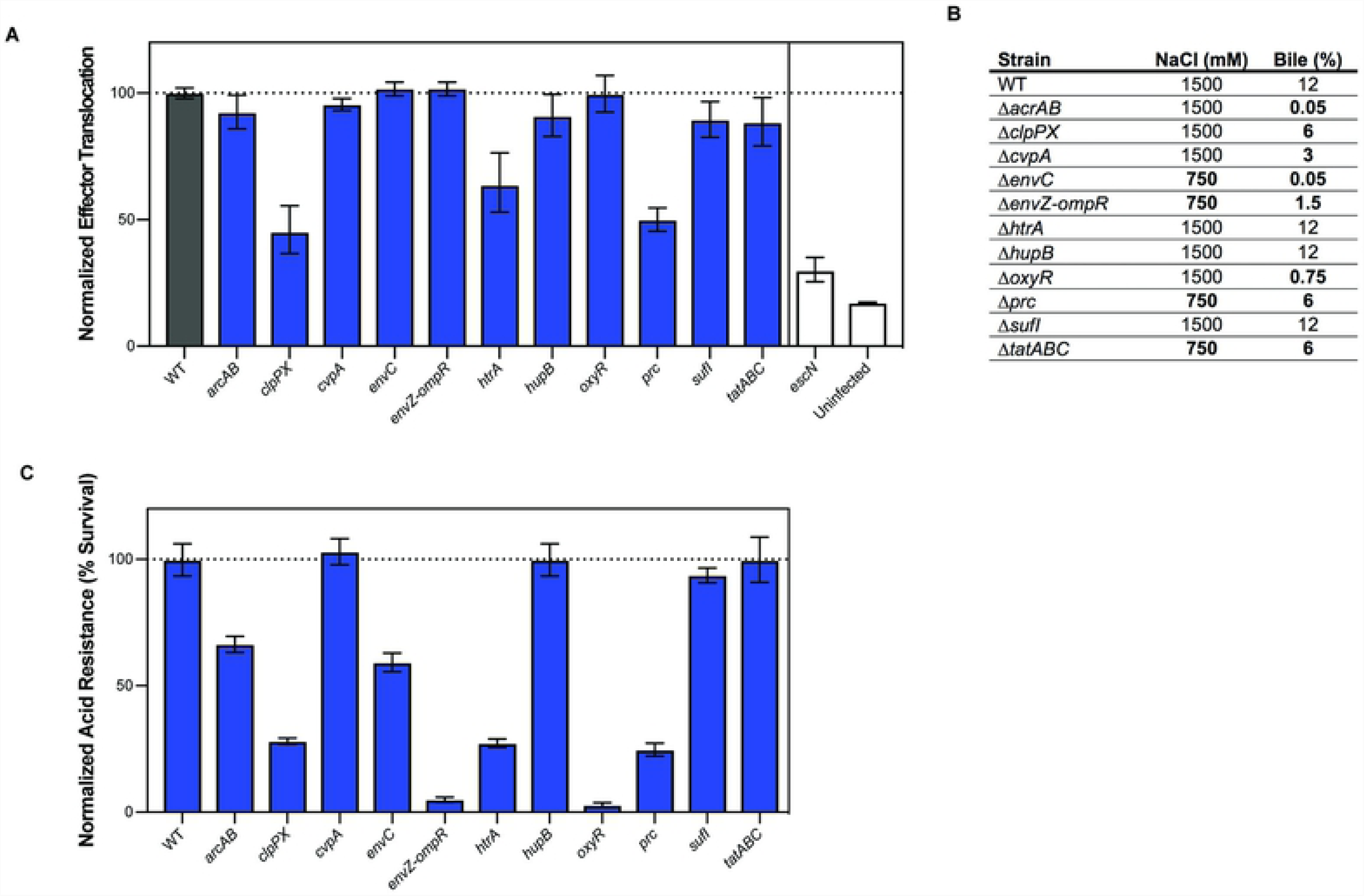
Effector translocation and survival in response to various gastro-intestinal stressors by mutants defective in colonization. A) Normalized effector translocation of mutants compared to WT. Δ*escN*, a mutant that abrogates T3SS activity, was used as a control. Mutants were tested for their ability to translocate EspF-TEM1 into HeLa cells, as measured by a shift in emission spectra from 520 to 450 nm. Fluorescence was normalized to WT levels. Geometric means and geometric standard deviations are plotted. B) MIC for NaCl (osmotic stress) and crude bile for the indicated mutants. Bold text highlights values differing from the wild-type. C) Normalized acid resistance. Mutants were tested for their ability to survive low acid shock. Survival is shown as a percentage of the acid resistance of the WT. Geometric mean and geometric standard deviation are plotted.

We also investigated the capacity of each of the 11 mutant strains to survive challenge with three stressors – low pH, bile, and high salt (osmotic challenge) – that the pathogen may encounter in the gastrointestinal tract. Relative to the WT strain, all but one (*sufI*) of the mutant strains exhibited reduced survival following one or more of these challenges (Fig 5BC), suggesting that exposure to these host environmental factors may contribute to the in vivo attenuation of these mutants. Many of the EHEC mutants exhibited sensitivities to external stressors that are consistent with previously described phenotypes in other organisms and experimental systems. For example, the EHEC Δ*acrAB* locus, which was associated with bile sensitivity in EHEC (Fig 5B), is known to contribute to a multidrug efflux system that can extrude bile salts, antibiotics, and detergents (84). Our observation that mutants lacking the oxidative stress response gene *oxyR* are sensitive to bile and to acid pH is also concordant with previous reports linking both stimuli to oxidative stress (85–87). Furthermore, the heightened sensitivity to bile, acid, and elevated osmolarity of EHEC lacking the two-component regulatory system EnvZ/OmpR is consistent with previous reports that EnvZ/OmpR is a critical determinant of membrane permeability, due to its regulation of outer membrane porins OmpF and OmpC. Mutations that activate this signaling system (in contrast to the deletions tested here) have been found to promote *E. coli* viability in vivo and to enhance resistance to bile salts (88).

The EHEC Δ*tatABC* mutant exhibited a marked colonization defect and a modest increase in bile sensitivity. The twin-arginine translocation (Tat) protein secretion system, which transports folded protein substrates across the cytoplasmic membrane (reviewed in (89,90)), has been implicated in the pathogenicity of a variety of Gram-negative pathogens, including enteric pathogens such as *Salmonella enterica* serovar Typhimurium (91–93), *Yersinia pseudotuberculosis* (94,95), *Campylobacter jejuni* (96), and *Vibrio cholerae* (97). Attenuation of Tat mutants can reflect the combined absence of a variety of secreted factors. For example, the virulence defect of *S. enterica* Typhimurium *tat* mutants are likely due to cell envelope defects caused by the inability to secrete the periplasmic cell division proteins AmiA, AmiC and SufI (92). Notably, single knock-outs of any of these genes did not cause attenuation (92), but altogether their absence renders the cell-envelope defective and more sensitive to cell-envelope stressors, such as bile acids (93).

In EHEC, the Tat system has been implicated in Stx1 export (98), but since Stx1 was not a hit in our screen and is not thought to modulate intestinal colonization (12), it is not likely to explain the marked colonization defect of the EHEC Δ*tatABC* mutant. The suite of EHEC Tat substrates has not been experimentally defined, although putative Tat substrates can be identified by a characteristic signal sequence (89,90). A few substrates, including SufI, OsmY, OppA, MglB, and H7 flagellin, have been detected experimentally (98). *sufI*, interestingly, was also a validated hit in our screen and is the only Con-ARTIST defined CD gene that has a predicted Tat-secretion signal. However, the Δ*sufI* mutant did not display enhanced bile sensitivity, suggesting that attenuation of this mutant, and perhaps of the Δ*tatABC* mutant as well, reflects deficiencies in other processes. SufI is a periplasmic cell division protein that localizes to the divisome and may be important for maintaining divisome assembly during stress conditions (99,100). *E. coli tat* mutants have septation defects (101), presumably from loss of SufI at the divisome. Interestingly, *envC*, another validated CD gene, encodes a septal murein hydrolase (102) that is required for cell division, and the Δ*envC* mutant also displayed increased bile sensitivity. Consistent with this hypothesis, in high osmolarity media, the Δ*sufI*, Δ*envC*, and Δ*tatABC* mutants exhibited septation or cell shape defects (Fig S4B). Collectively, these data suggest that an impaired capacity for cell division may reduce EHEC’s fitness for intraintestinal growth, and that at times this may reflect increased susceptibility to clearance by host factors such as bile.

### CvpA promotes EHEC resistance to deoxycholate

We further characterized EHEC Δ*cvpA* because other TIS-based studies of the requirements for colonization by diverse enteric pathogens (*Vibrio cholerae, Vibrio parahaemolyticus* and *Salmonella enterica* serovar Typhimurium) also classified *cvpA* as important for colonization, but did not explore the reasons for the colonization deficiency of the respective mutants (40,45,46).

*cvpA* encodes a putative inner membrane protein and has been linked to colicin V export in *E. coli* K-12 (103) as well as curli production and biofilm formation in UPEC (104). The EHEC Δ*cvpA* mutant did not exhibit an obvious defect in biofilm formation or curli production (Fig S5AB), suggesting that *cvpA* may have a distinct role in EHEC pathogenicity.

To further characterize the sensitivity of EHEC Δ*cvpA* mutant to bile, we exposed the mutant to the two major bile salts found in the gastrointestinal tract, cholate (CHO) and deoxycholate (DOC) (Fig 6AC) (85,105). Cholate is a primary bile salt, produced in the liver and released into the biliary tract, while deoxycholate is a secondary bile salt that is generated from cholate by intestinal bacteria in the colon. In contrast to WT EHEC, which displayed equivalent sensitivity to the two bile salts in MIC assays (MIC = 2.5% for both), the Δ*cvpA* mutant was much more sensitive to DOC than to CHO (MIC= 0.08% versus 1.25%). The Δ*cvpA* mutant’s sensitivity to deoxycholate was present both in liquid cultures and during growth on solid media (Fig. 6ABC).

**Figure 6:**
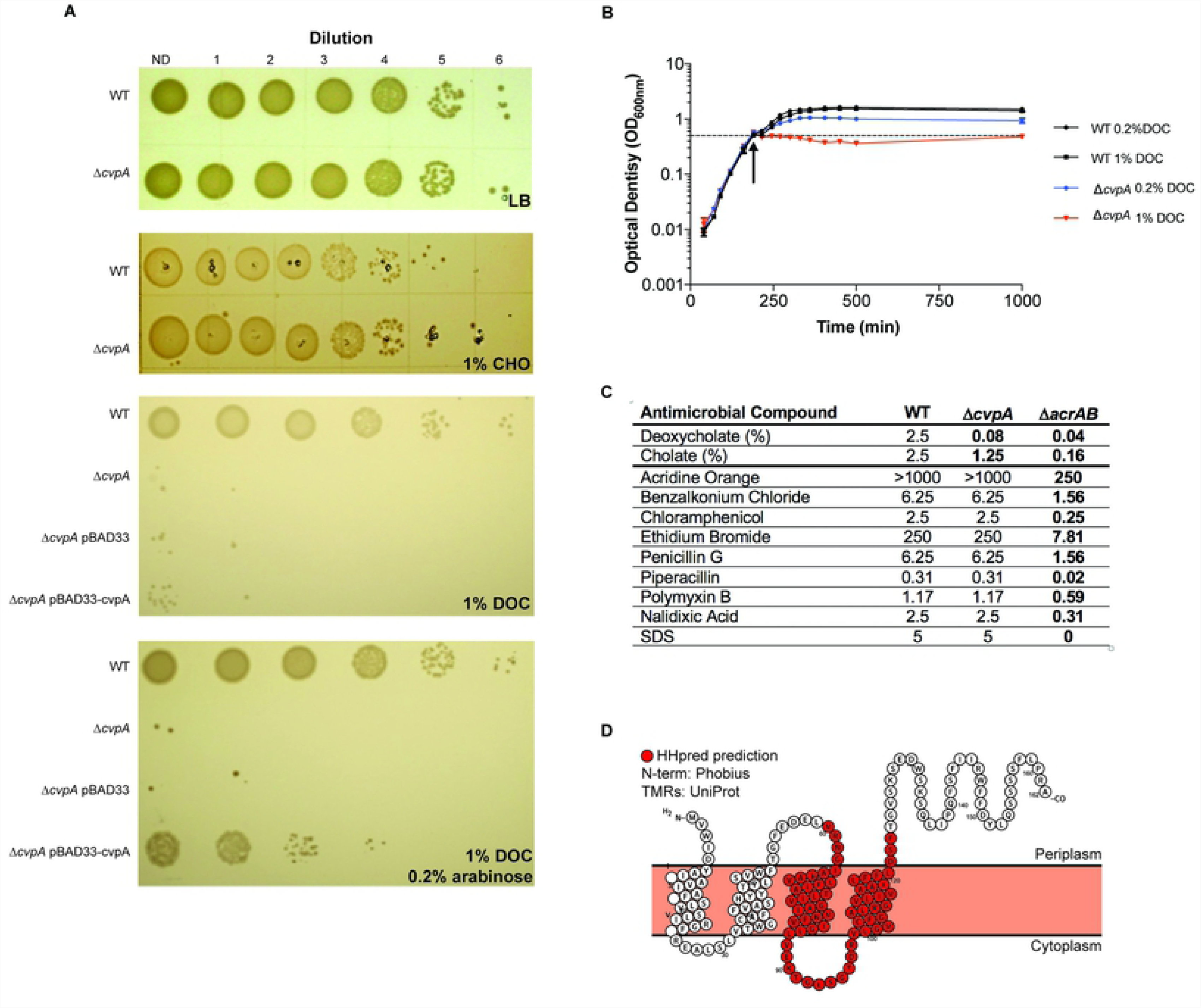
CvpA promotes EHEC resistance to deoxycholate. A) Dilution series of WT, Δ*cvpA* mutant, and Δ*cvpA* mutant with arabinose-inducible *cvpA* complementation plasmid plated on LB, LB 1% deoxycholate (DOC), LB 1% cholate (CHO), or LB 1% DOC + 0.2% arabinose. B) Optical density of WT and Δ*cvpA* grown in LB and two concentrations of DOC, added at the indicated arrow. The average of three readings is plotted with errors bars indicate standard deviation. C) MIC of antimicrobial compounds for WT and Δ*cvpA* and Δ*acrAB* mutants. Units are mg/mL unless specified otherwise. Bolded values are those different than the wild-type. D) Predicted CvpA topology diagram.

Growth of the Δ*cvpA* mutant in the presence of deoxycholate was partially restored by introduction of *cvpA* under the control of an inducible promoter, confirming that sensitivity is linked to the absence of *cvpA* (Fig. 6A)*. cvpA* lies upstream of the purine biosynthesis locus *purF*, and some Δ*cvpA* mutant phenotypes have been attributed to reduced expression on *purF* due to polar effects (103,106). The growth of the EHEC Δ*cvpA* mutant was not impaired in the absence of exogenous purines (Fig S5C), suggesting the *cvpA* deletion does not adversely modify *purF* expression.

Bile sensitivity has been associated with defects in the bacterial envelope or with reduced efflux capacity (reviewed in (105)). We assessed the growth of the Δ*cvpA* mutant in the presence of a variety of agents that perturb the cell envelope to assess the range of the defects associated with the absence of *cvpA*. The MICs of WT and Δ*cvpA* EHEC were compared to those of an Δ*acrAB* mutant, whose lack of a broad-spectrum efflux system provided a positive control for these assays. Notably, the Δ*cvpA* mutant did not exhibit enhanced sensitivity to any of the compounds tested other than bile salts. In marked contrast, the Δ*acrAB* mutant displayed increased sensitivity to all agents assayed (Fig 6C). These observations suggest that the sensitivity of the *cvpA* mutant to DOC is not likely attributable to a general cell envelope defect in this strain. *V. cholerae* and *V. parahaemolyticus* Δ*cvpA* mutants also exhibited sensitivity to DOC (Fig S5D), implying a similar role in bile resistance in these distantly related enteric pathogens.

A variety of bioinformatic algorithms (PSLPred, HHPred, Phobius, Phyre2) suggest that CvpA is an inner membrane protein with 4-5 transmembrane elements similar to small solute transporter proteins (Fig 6D). Phyre2 and HHPred reveal CvpA’s partial similarity to inner membrane transporters in the Major Facilitator Superfamily of transporters (MFS) and the small-conductance mechanosensitive channels family (MscC). Additional protein classification schemes group CvpA with proteins involved in solute transport. For example, the PFAM database groups CvpA (PF02674) in the LysE transporter superfamily (CL0292), a set of proteins known to enable solute export. In conjunction with findings presented above, these predictions raise the possibility that CvpA is important for the export of a limited set of substrates that includes DOC. Additional studies to confirm this hypothesis and to establish how CvpA enables export are warranted, particularly because this protein is widespread amongst enteric pathogens.

## Conclusions

Here, we created a highly saturated transposon library in EHEC EDL933 to identify the genes required for in vitro and in vivo growth of this important food-borne pathogen using TIS. This approach has transformed our capacity to rapidly and fairly comprehensively assess the contribution an organism’s genes to growth in different environments (41,107,108). However, technical and biologic issues can confound interpretation of genome-scale transposon-insertion profiles. For example, we found that EHEC genes with low GC content or those without homologs in K-12 were less likely to contain transposon-insertions (Fig S2BC, Fig1C). Many of these genes were likely acquired during EHEC evolution via lateral gene transfer; they constitute some of the ∼1.4MB of DNA that distinguishes EHEC from K-12 strains. Unexpectedly, more than 100 of the genes conserved between EHEC and K-12 appear to promote the growth of the pathogen in rich media but not that of K-12 (Table S5), suggesting that the ∼1.4MB of laterally acquired DNA that distinguishes EHEC and K-12 has enabled divergence of the metabolic roles of ancestral *E. coli* genes in these backgrounds.

In animal models of infection, bottlenecks that result in marked stochastic loss of transposon mutants can severely constrain TIS-based identification of genes required for in vivo growth. Analysis of the distributions of the EHEC transposon-insertions in vitro and in vivo (Fig. 2) revealed that there is a large infection bottleneck in the infant rabbit model of EHEC colonization. Both Con-ARTIST, which applies conservative parameters to define conditionally depleted genes (Fig 2B), and a PCA-based approach, CompTIS, were used to circumvent the analytical challenges posed by the severe EHEC infection bottleneck. These approaches should also be of use for similar bottlenecked data that often hampers interpretation of TIS-based infection studies. Validation studies, which showed that 14 of 17 genes (82%) classified as CD were attenuated for colonization, suggest that these approaches are useful. Besides the LEE-encoded T3SS, more than 200 additional genes were found to contribute to EHEC survival and/or growth within the intestine. This set of genes should be of considerable value for future studies elucidating the processes that enable the pathogen to proliferate in vivo and for design of new therapeutics.

## Materials and Methods

### Ethics statement

All animal experiments were conducted in accordance with the recommendations in the Guide for the Care and Use of Laboratory Animals of the National Institutes of Health and the Animal Welfare Act of the United States Department of Agriculture using protocols reviewed and approved by Brigham and Women’s Hospital Committee on Animals (Institutional Animal Care and Use Committee protocol number 2016N000334 and Animal Welfare Assurance of Compliance number A4752-01)

### Bacterial strains, plasmids and growth conditions

Strains, plasmids and primers used in this study are listed in Supplementary Tables 8 and 9. Strains were cultured in LB medium or on LB agar plates at 37°C unless otherwise specified. Antibiotics and supplements were used at the following concentrations: 20 µg/mL chloramphenicol (Cm), 50 µg/mL kanamycin (Km), 10 µg/mL gentamicin (Gent), 50 µg/mL carbenicillin (Carb), and 0.3 mM diaminopimelic acid (DAP).

A gentamicin-resistant mutant of *E. coli* O157:H7 EDL933 (Δ*lacI::aacC1*) and a chloramphenicol-resistant mutant of *E. coli* K-12 MG1655 (Δ*lacI::cat*) were used in this study for all experiments, and all mutations were constructed in these strain backgrounds except where specified otherwise. The Δ*lacI::aacC1* and Δ*lacI*::*cat* mutations were constructed by standard allelic exchange techniques (109) using a derivative of the suicide vector pCVD442 harboring a gentamicin resistance cassette amplified from strain TP997 (Addgene strain #13055) (110) or a chloramphenicol resistance cassette from plasmid pKD3 (Addgene plasmid #45604) (53) flanked by the 5’ and 3’ DNA regions of the *lacI* gene. Isogenic mutants of EDL933 Δ*lacI::aacC1* were also constructed by standard allelic exchange using derivatives of suicide vector pDM4 harboring DNA regions flanking the gene(s) targeted for deletion. *E. coli* MFDλpir (111) was used as the donor strain to deliver allelic exchange vectors into recipient strains by conjugation. Sequencing was used to confirm mutations.

A Δ*cvpA* strain was also constructed using standard allelic exchange in a streptomycin-resistant mutant (Sm^R^) of *V. parahaemolyticus* RIMD 2210633. A *cvpA*::tn mutant was used from a *Vibrio cholerae* C6706 arrayed transposon library (112).

### Transposon-insertion library construction

To create transposon-insertion mutant libraries in EHEC EDL933 Δ*lacI::aacC1*, conjugation was performed to transfer the transposon-containing suicide vector pSC189 (113) from a donor strain (*E. coli* MFDλpir) into the EDL933 recipient. Briefly, 100 µL of overnight cultures of donor and recipient were pelleted, washed with LB, and combined in 20 µL of LB. These conjugation mixtures were spotted onto a 0.45 µm HA filter (Millipore) on an LB agar plate and incubated at 37°C for 1 h. The filters were washed in 8 mL of LB and immediately spread across three 245×245 mm^2^ (Corning) LB-agar plates containing Gent and Kn. Plates were incubated at 37°C for 16 h and then individually scraped to collect colonies. Colonies were resuspended in LB and stored in 20% glycerol (v/v) at −80°C as three separate library stocks. The three libraries were pooled to perform essential genes analysis, and one library aliquot was used to as an inoculum for infant rabbit infection studies.

To create TIS mutant libraries in *E. coli* K-12 MG1655 Δ*lacI*::cat, conjugation was performed as above. 200 uL of overnight culture of the donor strain (*E. coli* MFDλpir carrying pSC189) and the recipient strain (MG1655 Δ*lacI*::cat) were pelleted, washed, combined and spotted on 0.45 µm HA filters at 37°C for 5.5 hours. Cells were collected from the filter, washed, plated on selective media (LB Kan, Cat), and incubated overnight at 30°C. Colonies were resuspended in LB and frozen in 20% glycerol (v/v). An aliquot was thawed and gDNA isolated for analysis.

### Infant rabbit infection with EHEC transposon-insertion library

Mixed gender litters of 2-day-old New Zealand White infant rabbits were co-housed with a lactating mother (Charles River). To prepare the EHEC transposon-insertion library for infection of infant rabbits, 1 mL from one library aliquot was thawed and added to 20 mL of LB. After growing the culture for 3 h at 37°C with shaking, the OD_600_ was measured and 40 units of culture at OD_600_=1 (about 8 mL) were pelleted and resuspended in 10 mL PBS. Dilutions of the inoculum were plated on LB agar plates with Gent and Km for precise dose determination. An aliquot of the inoculum was saved for subsequent gDNA extraction and sequencing (input). Each infant rabbit was infected orogastrically with 500 µl of the inoculum (1×10^9^ cfu) using a size 4 French catheter. Following inoculation, the infant rabbits were monitored at least 2x/day for signs of illness and euthanized 2 days postinfection. The entire intestinal tract was removed from euthanatized rabbits, and sections of the mid-colon were removed and homogenized in 1 mL of sterile PBS using a minibeadbeater-16 (BioSpec Products, Inc.). 200 uL of tissue homogenate from the colon were plated on LB agar + Gm + Km to recover viable transposon-insertion mutants. Plates were grown for 16 h at 37°C. The next day, colonies were scraped and resuspended in PBS. A 5 mL aliquot of cells was used for genomic DNA extraction and subsequent sequencing (Rabbits 1-7).

### Characterization of transposon-insertion libraries

Transposon-insertion libraries were characterized as described previously. Briefly, for each library, gDNA was isolated using the Wizard Genomic DNA extraction kit (Promega). gDNA was then fragmented to 400-600 bp by sonication (Covaris E220) and end repaired (Quick Blunting Kit, NEB). Transposon junctions were amplified from gDNA by PCR. PCR products were gel purified to isolate 200-500bp fragments. To estimate input and ensure equal multiplexing in downstream sequencing, purified PCR products were subjected to qPCR using primers against the Illumina P5 and P7 hybridization sequence. Equimolar DNA fragments for each library were combined and sequenced with a MiSeq.

Reads were first trimmed of transposon and adaptor sequences using CLC Genomics Workbench (QIAGEN) and then mapped to *Escherichia coli* O157:H7 strain EDL933 (NCBI Accession Numbers: chromosome, NZ_CP008957.1; pO157 plasmid, NZ_CP008958.1) using Bowtie without allowing mismatches. Reads were discarded if they did not align to any TA sites, and reads that mapped to multiple TA sites were randomly distributed between the multiple sites. After mapping, sensitivity analysis was performed on each library to ensure adequate sequencing depth by sub-sampling reads and assessing how many unique transposon mutants were detected (Fig S2). Next, the data was normalized for chromosomal replication biases and differences in sequencing depth using a LOESS correction of 100,000-bp and 10,000-bp windows for the chromosome and plasmid, respectively. The number of reads at each TA site was tallied and binned by gene and the percentage of disrupted TA sites was calculated. Genes were binned by percentage of TA sites disrupted (Fig 1A, 1C).

For essential gene analysis, EL-ARTIST was used as in (45). Protein-coding genes, RNA-coding genes, and pseudogenes were included in this analysis. Briefly, EL-ARTIST classifies genes into one of three categories (underrepresented, regional, or neutral), based on their transposon-insertion profile. Classifications are obtained using a hidden Markov model (HMM) analysis following sliding window (SW) training (p <0.05, 10 TA sites). Insertion-profiles for example genes were visualized with Artemis.

For identification of mutants conditionally depleted in the rabbit colon as compared to the input inoculum, Con-ARTIST was used as in (114). First, the input library was normalized to simulate the severity of the bottleneck as observed in the libraries recovered from rabbit colons using multinomial distribution-based random sampling (n=100). Next, a modified version of the Mann-Whitney U (MWU) function was applied to compare these 100 simulated control data sets to the libraries recovered from the rabbit colon. All genes were analyzed, but classification as “conditionally depleted” was restricted to genes that had sufficient data (≥5 informative TA sites), met our standard of attenuation (mean log_2_ fold change ≤ −2), met our standard of phenotypic consistency (MWU p-value of ≤0.05), and had a consensus classification in 5 or more of the 7 animals analyzed. Genes with ≥5 informative TA sites that fail to exceed both standards of attenuation and consistency are classified as “queried” (blue), whereas genes with less than 5 informative TA sites are classified as “insufficient data”.

Gene-level PCA (glPCA) was performed using CompTIS, a principal component analysis-based TIS pipeline, as described in (66). Briefly, log_2_ fold change values were derived by comparing read abundance in each sample to 100 control-simulated datasets as in Con-ARTIST. These fold change values were weighted to minimize noise due to variability (for details, see (66)). Next, genes that did not have a fold change reported for all 7 animals were discarded. The fold change values were then z-score normalized. Weighted PCA was performed in Matlab (Mathworks) with the PCA algorithm (pca).

### GC content

The GC content of classified genes was compared using a Mann-Whitney U statistical test and a Bonferroni correction for multiple hypothesis correction when more than one comparison was made. A p-value <0.05 was considered significant for one comparison, p<0.025 for two. A Fisher’s exact two-tailed t-test was used to compare ratios of classifications between groups, where a p-value of <0.01 was considered significant.

### In vivo competitive infection

Barcodes were introduced into Δ*lacI::aacC1*and isogenic mutant strains as described previously (45,115) (46,63) Briefly, a 991bp fragment of *cynX* (RS02015) that included 51bp of the intergenic region between *cynX* and *lacA* (RS02020) was amplified using primers that contained a 30 bp stretch of random sequence and cloned into SacI and XbaI digested pGP704. The resulting pSoA176.mix was transformed into *E. coli* MFDλpir. Individual colonies carrying unique tag sequences were isolated and used as donors to deliver pSoA176 barcoded derivatives to EDL933 Δ*lacI::aacC1*and each isogenic mutant strain. Three barcodes were independently integrated into EDL933 Δ*lacI::aacC1*, and three barcodes into each isogenic mutant via homologous recombination in the intergenic region between *cynX* and *lacA*, which tolerates transposon-insertion in vitro and in vivo, indicating this locus is neutral for the fitness of the bacteria. Correct insertion of barcodes was confirmed by PCR and sequencing.

To prepare the culture of mixed EHEC-barcoded strains for the multi-coinfection experiment, 100 µl of overnight cultures of the barcoded strains were mixed in a flask and 1 mL of this mix was added to 20 mL LB. After growing the culture for 3 h at 37°C with shaking, the OD_600_ was measured and 40 units of culture at OD_600_=1 (about 8 mL) were pelleted and resuspended in 10 mL PBS. Dilutions of the inoculum were plated in LB agar plates with Gent and Carb for precise cfu determination. 10 infant rabbits were inoculated and monitored as described above, and colon samples collected. Tissue homogenate was plated, and CFU were collected the following day. gDNA was extracted and prepared for sequencing as in (115).

The quantification of sequence tags was done as described by (115). In brief, sequence tags were amplified from the inoculum culture and libraries recovered from rabbit colons. The relative in vivo fitness of each mutant was assessed by calculating the competitive index (CI) as follows.

We compare two strains (Δ*lacI::aacC1* and isogenic mutant) in a population with frequencies *f*_*wt*_ and *f*_*mut,x*_ respectively where x is one of 17 mutant strains with a deletion in gene x. For simplicity, we assume here that both expand exponentially from a time point t_0_ to a sampling time point t_s_, their relative fitness (offspring/generation) is proportional to the competitive index CI: 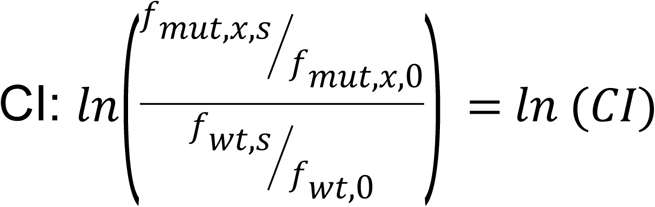. Here, *f*_*wt,0*_ and *f*_*mut,x,0*_ are the frequencies of the strains in the inoculum, measured in triplicates, and *f*_*wt,s*_and *f*_*mut,x,s*_ describe the frequencies at the sampling time point in the animal host. Because the WT strain was tagged with 3 individual tags and the inoculum was measured in triplicate, we have 3×3=9 measurements of the ratio *f*_*wt,s*_/ *f*_*wt*,0_. The same is true for all mutant strains, such that we have 9 measurements of the ratio *f*_*mut,x,s*_/*f*_*mut,x*,0_. In total, we therefore have 3×3×3×3= 81 CI measurements for each mutant per animal. To determine intra-host variance in these 81 measurements, a 95% confidence interval of the CI in single animal hosts was determined by bootstrapping. For combining the CIs measured across all 10 animal hosts, we performed a random-effects meta-analysis using the metafor package (116) in the statistical software package R (version 3.0.2). The pooled rate proportions and 95% confidence intervals were calculated using the estimates and the variance of CIs in each animal determined by bootstrapping and corrected for multiple testing using the Benjamini-Hochberg procedure.

### In vitro growth

Each bacterial strain was grown at 37°C overnight. The next day, cultures were diluted 1:1000 into 100 uL of LB in 96-well growth curve plates in triplicate. Plates were left shaking at 37°C for 10-24 hours. Absorbance readings at 600nm were normalized to a blank, and the average of each triplicate was taken as the optical density.

### T3SS translocation assays

T3SS functionality was assessed by translocation of the known EHEC T3SS effector protein EspF into HeLa cells as described previously (77). Briefly, the plasmid encoding the effector protein EspF fused to TEM-1 beta-lactamase was transformed into each of the bacterial strains to be tested. Overnight cultures of each bacterial strain were diluted 1/50 in DMEM supplemented with HEPES (25mM), 10% FBS and L-glutamine (2mM) and incubated statically at 37°C with 5% CO2 for two hours. This media is known to induce T3SS expression (117). HeLa cells were seeded at a density of 2×10^4^ cells in 96-well clear bottom black plates and infected for 30 minutes at an MOI of 100. After 30 minutes of infection IPTG was added at a final concentration of 1mM to induce the plasmid-encoded T3SS effector. After an additional hour of incubation, monolayers were washed in HBSS solution and loaded with fluorescent substrate CCF2/AM solution (Invitrogen) as recommended by the manufacturer. After 90 minutes, fluorescence was quantified in a plate fluorescence reader with excitation at 410nm and emission was detected at 450nm. Translocation was expressed as the emission ratio at 450/520nm to normalize beta-lactamase activity to cell loading and the number of cells presented at each well, and then normalized to WT levels of translocation.

### Biofilm, curli production, and purine assays

Biofilm and curli production assays were performed as described previously (104). For biofilm assays, bacterial cultures were grown in yeast extract-Casamino Acids (YESCA) medium until they reached an OD_600_ ∼ 0.5 and 1/1000 dilution of this culture was used to seed 96-well PVC plates. The cultures were grown at 30°C for 48 hours and biofilm production was quantitatively measured using crystal violet staining and absorbance reading at 595nm. Relative biofilm production was normalized to the average of three WT samples. To test curli production, bacterial cultures were grown in YESCA medium until they reached an OD_600_ ∼ 0.5 and then were struck to single colonies onto YESCA agar plates supplemented with Congo Red. Red colonies indicate curli production. To test if our Δ*cvpA* deletion had polar effects on *purF*, the mutant and WT were struck onto minimal media lacking exogenous purines.

### Acid shock assays

An adaptation of the acid shock method described in (118) was performed. Briefly, bacterial cultures were grown until mid-exponential phase (OD_600_ ∼ 0.6), then diluted 20-fold in LB pH 5.5 and incubated for 1 hour before preparing serial dilutions and plating each culture to determine the relative percentage of survival in comparison to the wild-type EDL933 strain. The pH of the LB broth was adjusted using sterilized 1mM HCl and buffered with 10% MES. Values are expressed as percent survival normalized to WT.

### MIC assays

MIC assays were performed using an adaptation of a standard methodology with exponential-phase cultures (119). Briefly, the different compounds to be tested (see Fig5B, 6B) were prepared in serial 2-fold dilutions in 50 ul of LB in broth in a 96-well plate format. To each well was added 50 ul of a culture prepared by diluting an overnight culture 1,000-fold into fresh LB broth, growing it for 1 h at 37°C, and again diluting it 1,000-fold into fresh medium. The plates were then incubated without shaking for 24 h at 37°C.

### Bile salts survival assays

Bile salt sensitivity assays were adapted from (120). For plate sensitivity assays, each bacterial strain was grown at 37°C until they reached mid-exponential phase of growth (OD_600nm_ of 0.5) and the culture was serially diluted and spot-titered onto LB agar plates supplemented with either 1% DOC or 1% CHO. Spots were air dried and plates incubated at 37°C for 24 h. For complementation, strains were grown in media and on plates supplemented with 0.2% arabinose. For sensitivity assays done in liquid culture, each bacterial strain was grown at 37°C until it reached mid-exponential phase of growth (OD_600nm_ of 0.5) and then cultures were split and supplemented with either DOC, CHO or buffer (PBS) and bacterial growth was assessed by absorbance at 600nm.

### Growth in high-salt media

Bacterial strains were grown in either LB or LB supplemented with 0.3M NaCl until mid-exponential phase and analyzed by phase microscopy.

### Computational Analysis

To enable comprehensive functional/pathway analyses in EHEC we carried out BLAST-based comparisons between the old EHEC genome sequence and annotation system (NCBI Accession Numbers AE005174 and AF074613) and the new sequence and annotation system (NZ_CP008957.1 and NZ_CP008958.1) (Table S1). This comparison links the new annotations (RS locus tags) to the original ‘Z numbers’ from (7) and their associated function and pathway annotation.

To make the correspondence table (Table S1) between the old EHEC annotation system (Z Numbers) and the new system (RS Numbers), local BLAST was used. First, a reference nucleotide database was generated from the newest EHEC sequence and annotation (NZ_CP008957.1 and NZ_CP008958.1). The EHEC genome sequence containing Z number annotations (AE005174 and AF074613) was used as the query. Best matches were taken as equivalent loci.

To find the K-12 homolog for EHEC genes (Table S2, column M), local BLAST was also used. A reference nucleotide and amino acid database was generated from MG1655 K-12 (NC_000913.3), and the newest EHEC genome sequence was used as the query. For pseudogenes and genes coding for RNA, ≥90% nucleotide identity across ≥90% of the gene length was considered a homolog. For protein coding genes, ≥90% amino acid identity across ≥90% of the amino acid sequence was considered a homolog.

To find KEGG pathways and COG assignments for genes of interest, the Z correspondence table was used to look up the Z number of each gene. The Z number and corresponding functional information was searched on the EHEC KEGG database.

To determine if COGs were enriched in certain groups of genes (such as conditionally depleted genes), a COG enrichment index was calculated as in (51). The COG Enrichment Index is the percentage of the genes of a certain category (essential genes or CD genes) assigned to a specific COG divided by the percentage of genes in that COG in the entire genome. A two-tailed Fisher’s exact test was used to determine if this ratio was independent of grouping. A Bonferroni correction was applied for multiple hypothesis testing. A p-value of <0.002 was considered to be significant.

Sequencing saturation of TIS libraries was determined by randomly sampling 100,000 reads from each library and identifying the number of unique mutants in that pool. Libraries are sequenced to saturation when no new mutants are identified as additional reads are added. 2-4 million reads are sufficient to capture the depth of libraries used here.

Several protein prediction programs (PSLPred, HHPred, Phobius, Phyre2) (121–123) were used to analyze the CvpA amino acid sequence. Protter (124) was used to compile information from several of these searches and generate a topological diagram. PRED-TAT (125) was used to search for tat-secretion signals in the list of CD genes.

## Supplementary Figure Legends

**S1: Sequencing saturation of TIS libraries**

Reads were randomly sampled from each library and the percentage of TA sites disrupted in each randomly selected pool were plotted for the EDL933 library (A), MG1655 library (B), the inoculum library used to infect infant rabbits (C), and the libraries recovered from 7 rabbit colons (D-J).

**S2: Assessment of non-neutral EHEC EDL933 genes.**

A) Non-neutral genes (defined by EL-ARTIST as either regional or underrepresented) by Clusters of Orthologous Groups (COG) classification. COG enrichment index (displayed as log2 enrichment) is calculated as defined in (51) as the percentage of the CD genes assigned to a specific COG divided by the percentage of genes in that COG in the entire genome. A two-tailed Fisher’s exact test with a Bonferroni correction was used to test the null hypothesis that enrichment is independent of TIS status. p-values considered to be significant if <0.002. Single asterisks (*) indicates p-value <0.002, double asterisks (**) indicate p-value <0.001, and triple asterisks (***) indicate p-value <0.0001.

B) GC content (%) of EDL933 genes classified as either neutral (blue) or non-neutral (regional + underrepresented; red) by TIS. Distributions are compared using a Mann-Whitney U non-parametric test; (***) p-value of <0.0001.

C) GC content (%) of EDL933 genes classified as either having homologs in MG1655 (homolog) or lacking homologs (divergent). Distributions are compared using a Mann-Whitney U test; (***)p-value of <0.0001.

D) TA insertions across *kdsC* in EDL933 (left) and MG1655 (right).

**S3: Con-ARTIST and CompTIS classification of genes important for colonization** A-G) Distribution of percentage TA site disruption in libraries recovered from 7 rabbit colons. These distributions are overlaid with Con-ARTIST classification (queried, blue; CD (conditionally depleted), red) as described in Figure 2B.

H) CD genes were grouped by consensus across animals. Many genes are CD in only one rabbit; fewer are classified as CD across all 7 animals. A standard of consensus of 5 or more animals was chosen to determine the list of CD genes, indicated with an asterisk (*).

I) Variance explained by each gene-level (gl) principal component for glPCA performed across the 7 rabbit screens.

J) Gene-level principal component 1 (glPC1) coefficients for each rabbit dataset.

H) Heatmap of the log2 fold change for each gene with a glPC1 score that falls within the bottom 10% of the distribution. Each column represents genes from a separate rabbit replicate. Genes are ordered by PC1 score, lowest at the top of the heatmap and highest at the bottom.

**S4: In vitro growth and morphology of mutants.**

A) 17 mutant strains plus the wild-type were grown in LB and turbidity measured by optical density. The average of three readings with the standard deviation is plotted.

B) Cell-shape defects of Δ*sufI*, Δ*envC*, Δ*tatABC*, and Δ*prc* mutants in high osmolality media. Morphology in LB (top) or LB supplemented with 0.3M NaCl (bottom) is shown.

**S5: Characterization of Δ*cvpA*.**

A) Biofilm production in WT and Δ*cvpA* using crystal violet staining and absorption. Levels were normalized to a percent of the WT value; three samples were analyzed and the geometric means and geometric standard deviation are plotted. The differences between the two groups were not significant (n.s.) by Mann-Whitney U.

B) WT and Δ*cvpA* struck to single colonies on an agar plate made with YESCA media supplemented with Congo Red to detect curli fibers.

C) WT and Δ*cvpA* struck to single colonies on an agar plate containing minimal media with no exogenous purines.

D) Dilution series of *Vibrio cholerae* C6706 WT and *cvpA*::tn plated on LB and LB 1% deoxycholate (DOC).

E) Dilution series of *Vibrio parahaemolyticus* WT and Δ*cvpA* plated on LB and LB 1% deoxycholate (DOC).

### Supplementary Tables Captions

S1) RS to Z Annotation

S2) EHEC EL-ARTIST

S3) EHEC Non-Neutral KEGG

S4) K12 EL-ARTIST

S5) EHEC Unique Non-Neutral

S6) Con-ARTIST and CompTIS

S7) EHEC CD Genes KEGG

S8) Strains

S9) Oligos

